# Choice of PP1 catalytic subunit affects neither requirement for G-actin nor insensitivity to Sephin1 of PPP1R15A-regulated eIF2αP dephosphorylation

**DOI:** 10.1101/228809

**Authors:** Ana Crespillo-Casado, Zander Claes, Meng S. Choy, Wolfgang Peti, Mathieu Bollen, David Ron

**Affiliations:** Cambridge Institute for Medical Research, University of Cambridge, Cambridge CB2 0XY, United Kingdom; Department of Cellular and Molecular Medicine, KU Leuven, BE-3000 Leuven, Belgium; Department of Chemistry and Biochemistry, University of Arizona, Tucson, Arizona 85721-0041, United States

**Keywords:** Phosphoprotein Phosphatase 1 (PP1), enzyme inhibitor, eukaryotic initiation factor 2a (eIF2a), G‐actin, Integrated Stress Response, proteostasis, protein synthesis, Sephin1, Guanabenz

## Abstract

The integrated stress response (ISR) is regulated by kinases that phosphorylate the α subunit of translation initiation factor 2 and phosphatases that dephosphorylate it. Genetic and biochemical observations indicate that the eIF2α^P^-directed holophosphatase - a therapeutic target in diseases of protein misfolding - is comprised of a regulatory, PPP1R15, and a catalytic, Protein Phosphatase 1 (PP1) subunit. In mammals, there are two isoforms of the regulatory subunit, PPP1R15A and PPP1R15B, with overlapping roles in promoting the essential function of eIF2α^P^ dephosphorylation. However, conflicting reports have appeared regarding the requirement for an additional co-factor, G-actin, in enabling substrate-specific de-phosphorylation by PPP1R15-containing PP1 holoenzymes. An additional concern relates to the sensitivity of the PPP1R15A-containing PP1 holoenzyme to the [(ochlorobenzylidene)amino]guanidines (Sephin1 or Guanabenz), small molecule proteostasis modulators. It has been suggested that the source and method of purification of the PP1 catalytic subunit and the presence or absence of an N-terminal repeat-containing region in the PPP1R15A regulatory subunit might influence both the requirement for G-actin by the eIF2α^P^-directed holophosphatase and its sensitivity to inhibitors. Here we report that in the absence of G-actin, PPP1R15A regulatory subunits were unable to accelerate eIF2α^P^ dephosphorylation beyond that affected by a catalytic subunit alone, whether PPP1R15A regulatory subunit had or lacked the N-terminal repeat-containing region and whether paired with native PP1 purified from rabbit muscle, or recombinant PP1 expressed in and purified from bacteria. Furthermore, none of the PPP1R15A-containing PP1c holophosphatases were inhibited by Sephin1 or Guanabenz.

---

The integrated stress response (ISR) is a signal transduction pathway that couples diverse stressful conditions to the activation of a rectifying translational and transcriptional program that is implicated in biological processes ranging from memory formation to immunity and metabolism (reviewed in Ref. 1). The mammalian ISR and its yeast counterpart (the general control response) are initiated by the phosphorylation of the α subunit of translation initiation factor 2 (eIF2α) on serine 51 (2,3) and its activity is terminated by eIF2α^P^ dephosphorylation.

Two related regulatory proteins, PPP1R15A/GADD34 and PPP1R15B/CReP, encoded in mammals by *PPP1R15A* and *PPP1R15B*, direct the unspecific Protein Phosphatase 1 (PP1) to promote eIF2α^P^ dephosphorylation (4-7). PPP1R15A or PPP1R15B form a complex with PP1 via a conserved region of ~70 amino acids (PPP1R15A residues 555-624) located at their C-termini (5,8-11) (**Fig. 1*A***). This conserved C-terminal region of either PPP1R15 regulatory subunit is sufficient to promote eIF2α^P^ dephosphorylation and to inactivate the ISR (4,5,10,11). Indeed, Herpes viruses have exploited this activity and encode a small protein homologous to the C-terminus of PPP1R15 to reverse eIF2α phosphorylation, undoing a defensive strategy of infected cells (12).

**Figure 1.**
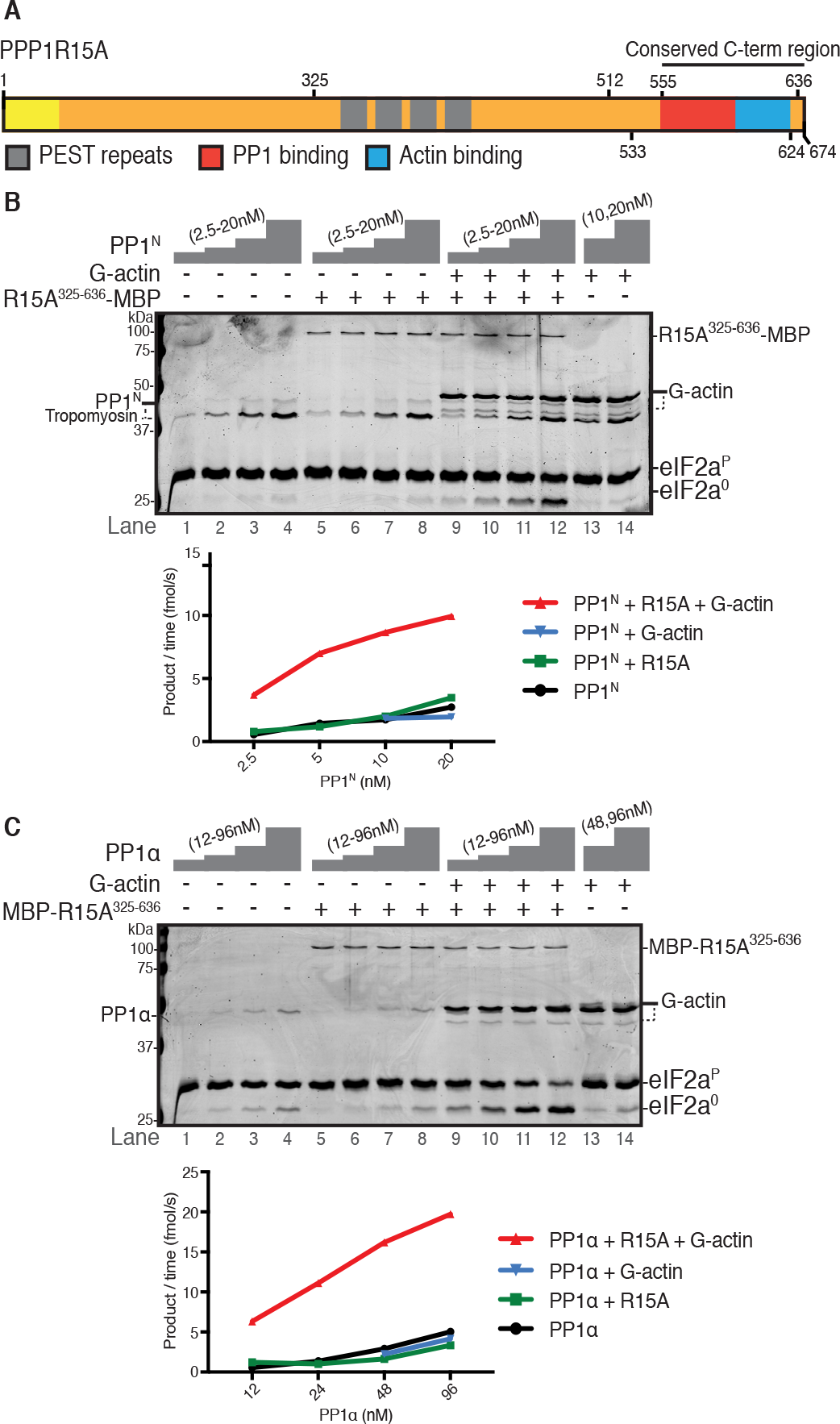
PPP1R15A accelerates eIF2α^P^ dephosphorylation by either PP1^N^ or PP1α only in the presence of G-actin. *A*, Cartoon representation of human PPP1R15A protein (1-674). Key residues used for truncated versions of the proteins in this study are annotated.The proline, glutamate, serine and threonine-rich (PEST) repeats are highlighted as are the PP1 and G-actin binding sites in the conserved C-terminal region. *B*, <u>Upper panel</u>: Coomassie-stained PhosTag-SDS-PAGE containing resolved samples of dephosphorylation reactions (30 minutes at 30˚C) in which 2 μM eIF2α^P^ was dephosphorylated by PP1^N^ purified from rabbit skeletal muscle in the presence or absence of PPP1R15A^325-636^-MBP (50 nM) and/or G-actin (400 nM). The position of the various protein species is indicated. eIF2α^P^ and eIF2α^0^ refer to the phosphorylated and non-phosphorylated form of the bacterially-expressed N-terminal domain (residues 1-185) of eIF2α, respectively. Note that both G-actin and PP1^N^ preparation gave rise to two bands: a major full-length species and minor degradation product, in the case of G-actin, and a PP1 and tropomyosin band in the case of PP1^N^ (see also Fig. *S1*). Shown is a representative experiment of four independent repetitions performed. <u>Lower panel</u>: Plot of the rate of eIF2α^P^ dephosphorylation as a function of the concentration of PP1^N^ in the experiment above. *C*, as in “*B*” but using bacterially-expressed PP1α as the catalytic subunit (96, 48, 24 or 12 nM), MBP-PPP1R15A^325-636^ (50 nM) and G-actin (400 nM). The assays were performed during 20 minutes at 30˚C. Shown is a representative experiment of three independent repetitions performed.

Despite genetic evidence pointing to the sufficiency of the conserved C-terminal portion of PPP1R15 in reversing the eIF2α^P^-dependent ISR *in vivo* (4,5,10), complexes formed *in vitro* between PPP1R15 regulatory subunit fragments and PP1 have been observed to unexpectedly lack specificity towards eIF2α^P^ (10). Dephosphorylation of eIF2α^P^ is no faster by a complex of PPP1R15A-PP1 (or PPP1R15B-PP1) than by PP1 alone, showing that when added as single components, PPP1R15A/B do not influence *k*_cat_ or *K*_m_ of PP1 towards the specific substrate eIF2α^P^ (10). However, addition of G-actin to the binary complex of PPP1R15 and PP1 selectively accelerates eIF2α^P^ dephosphorylation. G-actin binds directly to the conserved C-terminus of PPP1R15, alongside PP1 to form a ternary complex, whose affinity (*K_d_*~10^-8^ M) matches the EC_50_ of G-actin’s stimulatory effect (10,13). The *in vivo* relevance of G-actin to eIF2α^P^ dephosphorylation is attested to by the finding that actin sequestration in fibres (as F-actin) enfeebles eIF2α^P^ dephosphorylation, implying a role for factors that affect the actin cytoskeleton in ISR regulation (14).

The ability to dephosphorylate eIF2α^P^ is an essential function in developing mammals (15). Nonetheless, inactivation of the *PPP1R15A* gene, which decelerates eIF2α^P^ dephosphorylation and prolongs the ISR, is protective in certain cellular and animal models of diseases associated with enhanced unfolded protein stress (16-19). This has generated interest in targeting the PPP1R15A-containing holophosphatase for inhibition by small molecules (reviewed in Ref. 20), an endeavour that requires detailed knowledge of the enzymatic mode of action.

A recent report challenged the need for G-actin as a co-factor in PPP1R15A-mediated eIF2α^P^ dephosphorylation (21). Instead, it suggested that a binary complex assembled from PP1α and a fragment of PPP1R15A (PPP1R15A^325-636^), encompassing both the C-terminal PP1-binding region and the N-terminal repeat-containing extension, dephosphorylates eIF2α^P^ faster than PP1 alone (21). Importantly, dephosphorylation of eIF2α^P^ by this active binary complex was reported to be selectively inhibited *in vitro* by Guanabenz and Sephin1, two structurally-related small molecules that function *in vivo* as proteostasis modifiers (22,23). The new study contradicts previous observations that neither the non-selective PPP1R15A-PP1 binary complex, nor the eIF2α^P^-selective PPP1R15A-PP1-G-actin ternary complex, were susceptible to inhibition by Guanabenz or Sephin1 (9,13).

Here we address three important questions raised by these discrepant reports: Does the choice of PP1 catalytic subunit influence the requirement for G-actin by the eIF2α^P^–directed holophosphatase? What role does the N-terminal repeat-containing region of PPP1R15A play in eIF2α^P^ dephosphorylation by the holophosphatase? Do these factors influence the sensitivity of eIF2α^P^ dephosphorylation to Guanabenz and Sephin1?

## Results

### Both native PP1 and bacterially-expressed PP1α require the presence of G-actin to promote PPP1R15A-regulated eIF2α^P^ dephosphorylation

PP1 produced in *E. coli* may differ in its enzymatic activity from PP1 purified from animal tissues, both in its substrate specificity and in its sensitivity to regulatory subunits (reviewed in ref. 24). To determine if the G-actin-dependence of PP1-PPP1R15A-mediated eIF2α^P^ dephosphorylation is a peculiarity of the bacterially-expressed PP1γ isoform used previously (10,13), we purified the native catalytic subunit of PP1 from rabbit skeletal muscle (PP1^N^), following an established protocol (25), and compared the two PP1 preparations. Native PP1 (PP1^N^) is a mixture of PP1α, PP1β and PP1γ isoforms and gave rise to two prominent bands on SDS-PAGE (**Fig. S1*A***, left panel). The mass spectra of tryptic peptides derived from the PP1^N^ sample was analysed by Maxquant with iBAQ (intensity based absolute quant) to identify the major contaminating species (tropomyosin), and to estimate the relative contribution of PP1 and contaminants to the protein preparation. This enabled a comparison of the catalytic subunit content of PP1^N^ preparation with the bacterially-expressed PP1γ, which served as a reference.

Residues 1-324 of PPP1R15A mediate membrane association (26), but compromise solubility *in vitro*. Therefore, we used a PPP1R15A^325-636^ fragment lacking the membrane-anchoring domain, which is soluble when expressed in *E. coli*. **Fig. 1*B*** shows that addition of either PPP1R15A^325-636^, expressed and purified with a C-terminal maltose-binding protein (MBP) tag (lanes 5-8), or G-actin alone (lanes 13 & 14) did not stimulate eIF2α^P^ dephosphorylation by nanomolar concentrations of PP1^N^. However, addition of both G-actin and PPP1R15A^325-636^-MBP (lanes 9-12) stimulated dephosphorylation 5-fold (**Fig. 1*B***), similar to the increase observed with bacterially expressed PP1γ (**Fig. S1*B***)(10).

PP1 purified from rabbit muscle is a mixture of α, β and γ isoforms, whereas it has been reported that the PP1α isoform possesses *in vivo* selectivity for PPP1R15A (6). Therefore, we prepared bacterially-expressed PP1α by a method that promotes its native-like state (27). To control for effects the location of the tag might have on activity, we also generated an N-terminally MBP-tagged PPP1R15A^325-636^. The holophosphatase comprised of PP1α and MBP-PPP1R15A^325-636^ also exhibited a stringent requirement for G-actin (**Fig. 1*C***).

A concentration-dependent stimulatory effect of PPP1R15A on eIF2α^P^ dephosphorylation by the three component holoenzyme (PP1, PP1R15A and G-actin) was observed with constructs tagged at either their N- or C-termini and with either native or bacterially-expressed PP1 (**Fig. 2*A*** **and *B***). The difference in EC_50_ values obtained for PPP1R15A^325-636^-MPB with PP1^N^ (23 nM) or MBP-PPP1R15A^325-636^ with PP1α (6 nM) may reflect the effect of the position of the MBP-tag, the contaminating tropomyosin (in PP1^N^), or both. Importantly, the data agreed with similar experiments in which PPP1R15A^325-636^ and bacterially-expressed PP1γ were used, EC_50_ of 10 nM (Ref. 13, figure 8A therein).

**Figure 2.**
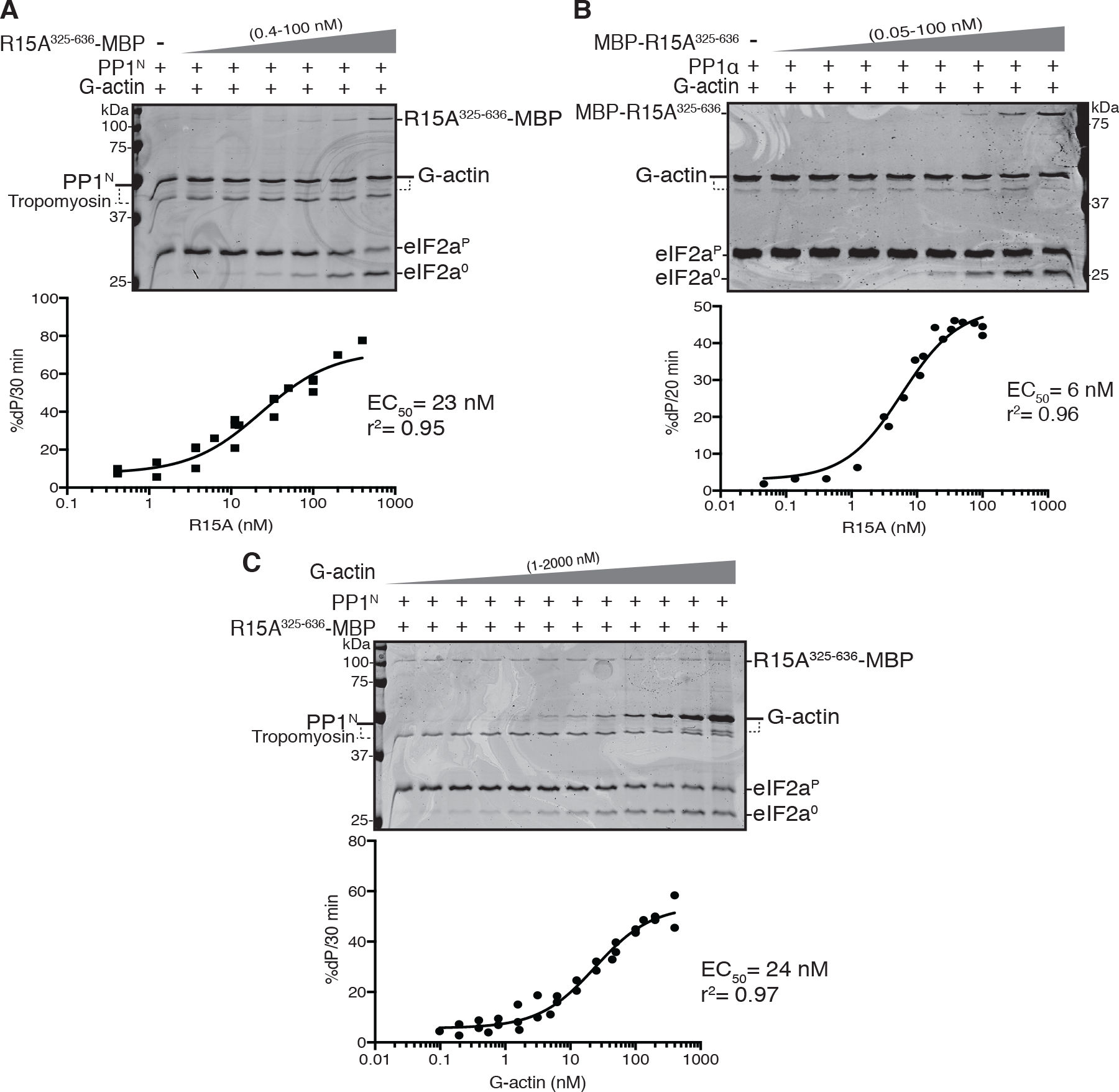
The source of catalytic subunit does not affect the kinetics of PPP1R15A and G-actin-mediated stimulation of eIF2α^P^ dephosphorylation. *A*, <u>Upper panel</u>. Coomassie-stained PhosTag-SDS-PAGE of dephosphorylation reactions (30 minutes at 30˚C), in which 2 μM eIF2α^P^ was dephosphorylated by PP1^N^ (20 nM) in presence of G-actin (400 nM) and increasing concentrations of PPP1R15A^325-636^-MBP (0-100 nM). Shown is a representative experiment of three independent experiments performed. <u>Lower panel</u>. Plot of the rate of dephosphorylation of eIF2α^P^ as a function of PPP1R15A^325-636^-MBP concentration, from the three experiments performed. The EC_50_ was calculated using the ”[Agonist] vs. response (three parameters)” function in GraphPad Prism v7. *B*, as in “*A*” but using bacterially-expressed PP1α (24 nM) and increasing concentrations of MBP-PPP1R15A^325-636^ (0-100 nM) in reactions performed over 20 minutes at 30˚C. Shown is a representative experiment of three independent experiments performed. *C*. As in “*A*” but with fixed concentrations of PP1^N^ (20 nM) and PPP1R15A^325-636^-MBP (50 nM) and varying the concentrations of G-actin (1-2000 nM). Shown is a representative experiment of three independent experiments performed.

G-actin also exerted a saturateable concentration-dependent stimulatory effect on the activity of a three-component holophosphatase constituted with native PP1^N^ **(Fig. 2*C*)**. The EC_50_ for G-actin with PP1^N^ (24 nM) was similar to that previously observed using bacterially-expressed PP1γ, EC_50_ of 13 nM (Ref. 13, figure 2C therein). Hence, despite variations in the estimated EC_50_ values for PP1R15A or G-actin, the combinations of catalytic and regulatory subunits tested showed consistent PPP1R15A and G-actin concentration dependent enzymatic activity. These experiments, conducted over a physiological concentration range (nanomolar catalytic subunit and micromolar substrate) and over a timescale aimed to minimize the effect of substrate depletion on enzyme kinetics, indicate that neither the source of PP1 nor the position of the tag in PPP1R15A are likely to account for the reported G-actin independent ability of PPP1R15A to stimulate eIF2α^P^ dephosphorylation.

### Lengthy incubation of the enzymatic reactions does not uncover PPP1R15A’s ability to promote G-actin-independent eIF2α^P^ dephosphorylation

Upon inhibition of the phosphorylating kinase, the eIF2α^P^ signal decays with a T_1/2_ of <10 minutes (with no change in the total eIF2α content) in both cultured mouse fibroblasts (Ref. 14, figure 6 therein) and Chinese Hamster Ovary cells (Ref. 13, figure 10 therein). Despite the rapid *in vivo* kinetics of the dephosphorylation reaction, the experiments pointing to G-actin-independent eIF2α^P^ dephosphorylation were conducted with long incubations of 16 hours at 30˚C (21). In the absence of other components PP1α is markedly unstable at 30˚C, losing about half of its activity by 1 hour and all detectable activity by 3 hours (**Fig. S2*A*** **and *B***). Thus a stabilizing effect of a PP1 binding co-factor might have accounted for the apparent G-actin-independent stimulatory effect of MBP-PPP1R15A^325-636^ on PP1α-mediated eIF2α^P^ dephosphorylation. However, over a range of PP1 concentrations (0.2-200 nM), the presence of MBP-tagged PPP1R15A^325-636^ failed to stimulate eIF2α^P^ dephosphorylation, whether PP1^N^ (**Fig. 3*A***) or PP1α (**Fig. 3*B***) were used as the catalytic subunit.

**Figure 3.**
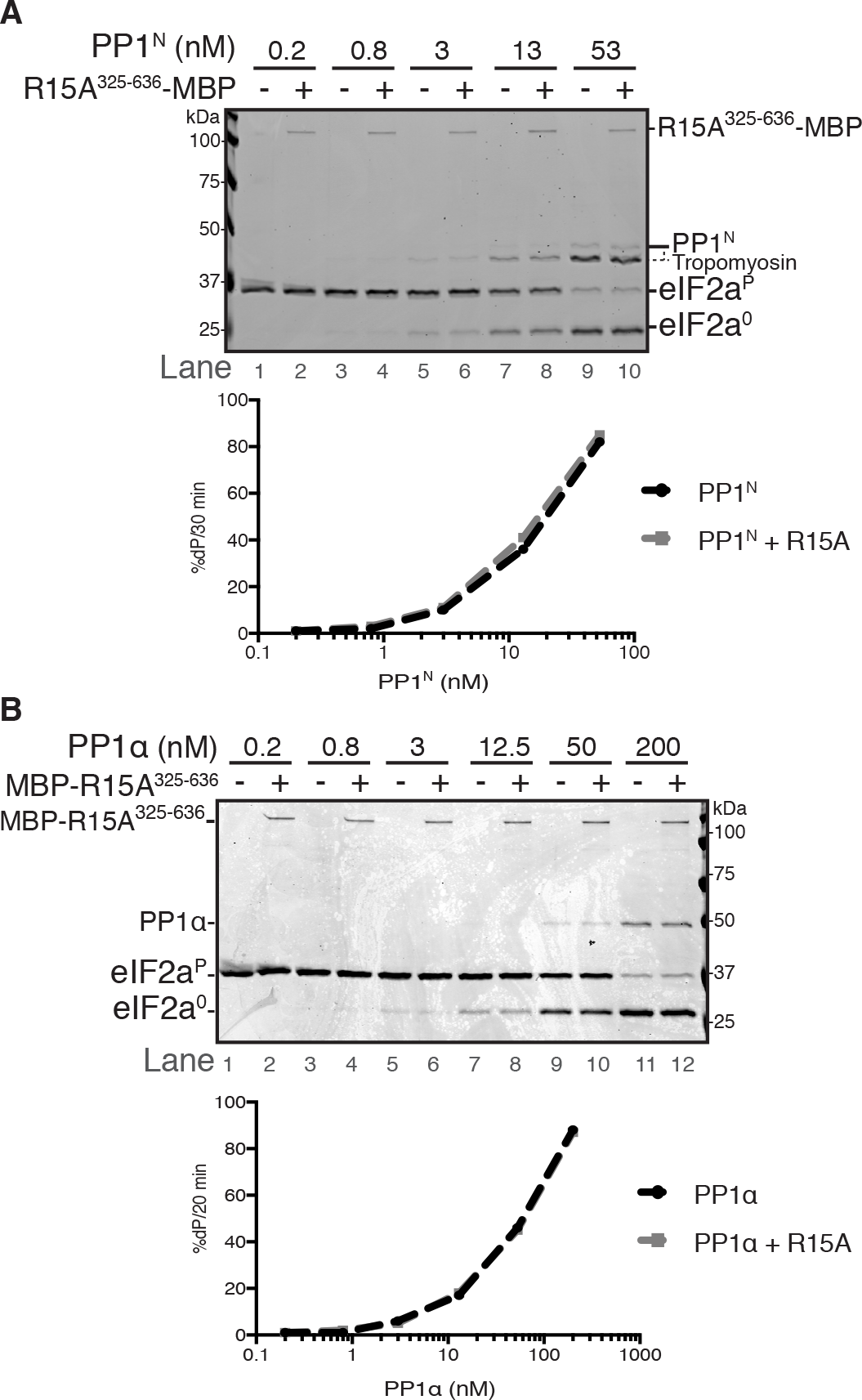
In the absence of G-actin, PPP1R15A is unable to stimulate dephosphorylation of eIF2αP (despite an extended incubation of 16h). *A*, <u>Upper panel</u>. Coomassie-stained PhosTag-SDS-PAGE containing dephosphorylation reactions (16 hours at 30˚C) in which 2 μM eIF2α^P^ was dephosphorylated by the indicated concentration of PP1^N^ in the presence or absence of PPP1R15A^325-636^-MBP (50 nM). Quantification of percentage of dephosphorylation (%dP) is shown below the image. Shown is a representative experiment of two independent repetitions performed.<u>Lower panel</u>. Plot of the rate of dephosphorylation of eIF2α^P^ as a function of PP1^N^ concentration. Data was obtained by quantification of bands of image shown above. *B*, as in “*A*” but using PP1α as the source of catalytic subunit and MBP-PPP1R15A^325-636^ (50 nM) as the regulatory subunit. Shown is a representative experiment of two independent repetitions performed.

### Substrate recruitment by the repeat-containing PPP1R15A^325-512^ region plays a secondary role in the kinetics of eIF2α^P^ dephosphorylation and its disruption is unlikely to account for sensitivity to Sephin1

PPP1R15A interacts directly with eIF2α, both in cells (9) and *in vitro* (21). This interaction maps to the repeat-containing region of PPP1R15A, residues 325-512; N-terminal to PPP1R15A’s PP1-binding domain (**Fig. 1*A***) and was proposed to play an important role in the catalytic cycle of PPP1R15A-containing holoenzymes (21). However, in the presence of G-actin, PPP1R15A^325-636^ and PPP1R15A^533-624^ stimulated eIF2α^P^ dephosphorylation similarly, when paired either with PP1^N^ (compare **Fig. 2*A*** and **4*A*** here) or with PP1γ (compare Figure 8A and Figure 2B in Ref. 13). These findings suggest that the conserved C-terminal PPP1R15 fragment that binds PP1 and G-actin simultaneously is sufficient to promote eIF2α^P^ dephosphorylation and to dominate its kinetics *in vitro* and call in to question the importance of the N-terminal repeats in PPP1R15A to the fundamentals of the holoenzyme’s catalytic cycle.

We considered that an important contributory role for substrate engagement by the PPP1R15A^325-533^ repeat-containing fragment to the catalytic cycle of the holophosphatase might have been masked by compensatory features that diverge between the different regulatory subunit constructs, fortuitously equalizing their activity. To address this possibility, we measured the ability of an MBP-PPP1R15A^325-512^ fragment (containing the repeats but lacking the C-terminal PP1 binding region) to compete with MBPPPP1R15A^325-636^-mediated (G-actin-dependent) eIF2α^P^ dephosphorylation using PP1α as the catalytic subunit. Minimal inhibition of the dephosphorylation reaction was observed at competitor concentrations of up to 8 µM (**Fig. 4*B***), which is a >300-fold excess over the MBPPPP1R15A^325-636^ regulatory subunit (present in the reaction at 24 nM), and a concentration of 18-fold above the reported *K*_d_ of the interaction between MBP-PPP1R15A^325-512^ and eIF2α^P^ (21). These data suggest that substrate recruitment by the N-terminal extension of PPP1R15A plays a secondary role in the kinetics of the dephosphorylation reaction and that the reported role of Sephin1 and Guanabenz in disrupting that interaction is unlikely to make an important contribution to their pharmacological activity. Consistent with these conclusions we find that under conditions in which eIF2α^P^ dephosphorylation is dependent on the concentration of both PP1α and MBPPPP1R15A^325-636^, we were unable to observe an inhibitory effect of Sephin1 or Guanabenz at a concentration of 100 µM (**Fig. 5**), which exceeds by two-fold that required for a proteostatic effect in cultured cells (13,23).

**Figure 4.**
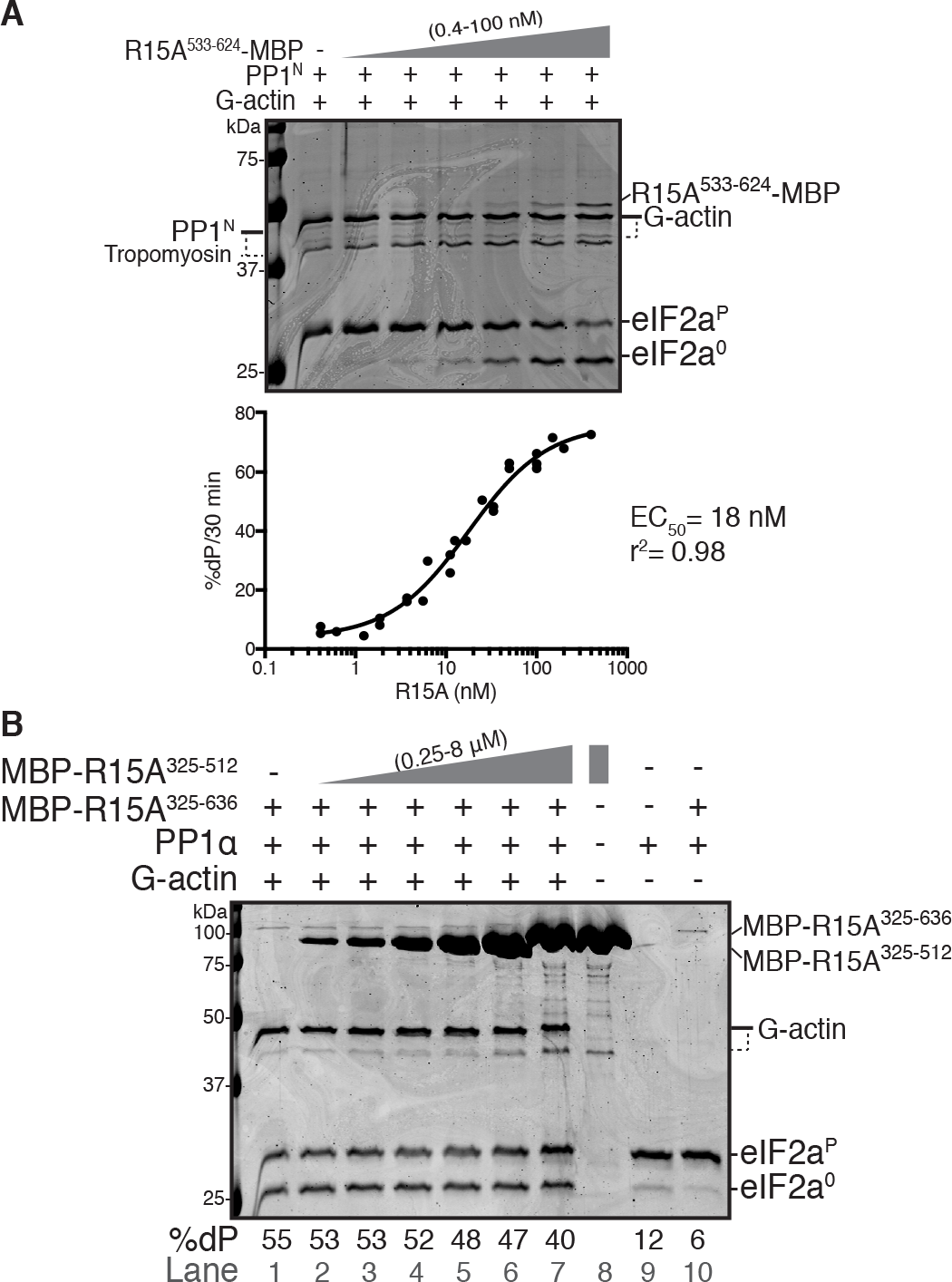
The C-terminal portion of PPP1R15A is sufficient to promote eIF2αP dephosphorylation. *A,* <u>Upper panel</u>: Coomassie-stained PhosTag-SDS-PAGE containing resolved samples from dephosphorylation reactions (30 minutes at 30˚C) in which 2 μM eIF2α^P^ was dephosphorylated by PP1^N^ (20 nM) in the presence of G-actin (400 nM) and increasing concentrations of PPP1R15A^533-624^-MBP (0-100 nM). Shown is a representative experiment of three independent repetitions performed. <u>Lower panel</u>: Plot of the rate of dephosphorylation of eIF2α^P^ as a function of PPP1R15A^533-624^-MBP concentration, from the three experiments performed. The EC_50_ was calculated using the ”[Agonist] vs. response (three parameters)” function in GraphPad Prism v7. *B*. as in “*A*” but using PP1α (24 nM) in presence of MBP-PPP1R15A^325-636^ (24 nM), G-actin (400 nM) and increasing concentrations of MBP-PPP1R15A^325-512^ as a competitor (0-8 μM). The assays were performed during 20 minutes at 30˚C. Lane 8, loaded with only MBP-PPP1R15A^325-512^ shows the absence of a species co-migrating with eIF2α^0^ (which might otherwise obscure an inhibitory effect on dephosphorylation). Lanes 9 and 10 control for the dependence of enzymatic activity on PPP1R15A and G-actin in this experiment. Quantification of percentage of dephosphorylation (%dP) is shown below the image. Shown is a representative experiment of two independent repetitions performed.

## Discussion

The new experiments presented here cover a range of conditions with realistic concentrations and time regimes. Incorporation of multiple time points and titrations of reaction components enabled a comparison of enzyme kinetics that accounts for the effect of substrate depletion. Our observations were made with two different PPP1R15A preparations and three different PP1 preparations, all of which consistently show the requirement for G-actin as an additional co-factor in enabling PPP1R15A to stimulate eIF2α^P^ dephosphorylation *in vitro*. As such the results presented here are in keeping with previous observations that G-actin has an essential role in promoting eIF2α^P^ dephosphorylation both *in vitro* and *in vivo* (10,13,14).

Our experiments also cast doubt on the importance of the physical interaction between the repeat-containing region of PPP1R15A (residues 325-512) and eIF2α^P^ in the substrate-specific dephosphorylation reaction. PPP1R15 regulatory subunits are found throughout the animal kingdom, yet only their C-terminal ~70 residues are conserved (11). This C-terminal fragment contains all the information needed to promote eIF2α^P^ dephosphorylation, as exemplified by its selective hijacking by Herpes viruses (12) and by experimentally targeted expression in cells (Ref. 10, figure 1C, therein). In complex with G-actin, the conserved C-terminal fragment of the PPP1R15s is also able to direct PP1 to selectively dephosphorylate eIF2α^P^ *in vitro* (Fig. 2 and 4 here and Ref. 10,13). None of this argues against an important regulatory role for the divergent N-terminal extensions of PPP1R15 regulatory subunits *in vivo*. This role may play out in terms of subcellular localization (26) or protein stability (28) and might be influenced by a physical interaction with the substrate (9,21). However, our findings argue that the physical interaction between PPP1R15A residues 325-512 and eIF2α^P^ is unlikely to play an important role in the formation of the enzyme-substrate complex required for catalysis and hence its disruption by Guanabenz or Sephin1 is unlikely to underscore any inhibitory effect they might have on eIF2α^P^ dephosphorylation

Most importantly perhaps, the findings presented here argue that the inability of previous efforts to uncover a role for Guanabenz or Sephin1 in inhibiting eIF2α^P^ dephosphorylation *in vitro* (9,13) was unlikely to have arisen from choice of catalytic subunit or from features of the PPP1R15A regulatory subunit used. Rather, the findings reported here, made *in vitro*, reinforce observations that Sephin1 and Guanabenz have no measurable effect on the rate of eIF2α^P^ dephosphorylation in cells (13). The recent description of PPP1R15A-independent cellular effects of Guanabenz (29) and our observations of Sephin1-induced proteostatic changes in gene expression, both in cells lacking PPP1R15A and in ISR-deficient cells with non-phosphorylatable eIF2α (13), suggest the need to revisit the role of these two compounds as eIF2α^P^ dephosphorylation inhibitors.

## Experimental procedures

### Protein expression and purification

The plasmids used to express protein in *E. coli* are presented in **Table S1**.

<u>PPP1R15A^325-636^-MBP and PPP1R15A^533-624^-MBP</u> were purified according to (13). Briefly, proteins were expressed in *E. Coli* BL21 (C3013) as N-terminally-tagged glutathione S-transferase fusion protein and purified by tandem affinity chromatography. First by binding to a glutathione sepharose 4B resin, elution with glutathione, followed by an overnight cleavage with Tobacco Etch Virus (TEV) protease (to remove the GST tag), binding to amylose beads and elution in maltose-containing buffer.

<u>MBP-PPP1R15A^325-636^ and MBP-PPP1R15A^325-512^</u>: *E. coli* BL21 (C3013) were transformed with the ampicillin-resistance pMAL-c5x-His vector (New England Biolabs, Cat. No. N8114) containing PPP1R15A^325-636^ and were selected in LB agar plates supplemented with 100 μg/ml ampicillin. A single colony was picked to grow overnight in 5 mL starter culture that served to inoculate 2 L of LB media (all supplemented with 100 μg/mL ampicillin), which was kept at 37°C. At OD_600_=0.6-0.8 protein expression was induced using 1 mM Isopropyl β-D-1-thiogalactopyranoside (IPTG) at 18°C for 20 hours. Bacteria were pelleted and resuspended in ice-cold His6 Lysis Buffer containing 50 mM Tris pH 7.4, 500 mM NaCl, 1 mM MgCl_2_, 1 mM tris (2-carboxyethyl)phosphine (TCEP), 100 μM phenylmethylsulfonyl fluoride (PMSF), 20 mTIU/ml aprotinin, 2 μM leupeptin, and 2 μg/ml pepstatin 20 mM imidazole and 10% glycerol. Bacterial suspensions were lysed using an Emulsi-Flex-C3 homogenizer (Avestin, Inc, Ottawa, Ontario) and clarified in a JA-25.50 rotor (Beckman Coulter) at 33,000 x g for 30 min at 4°C. Pre-equilibrated Ni-NTA beads (Qiagen, Cat. No. 30230,) were incubated with the samples for 2 hours at 4°C. Proteins were eluted in 2 mL of Imidazole Elution Buffer (50 mM Tris, pH 8, 100 mM NaCl, 500 mM Imidazole, 10% glycerol) and incubated with amylose beads (New England Biolabs, Cat No. E8021S) pre-equilibrated with Lysis Buffer (His6 Lysis Buffer without imidazole) for 2 hours at 4°C. The amylose beads were batch-washed using 25 bed volumes of Lysis Buffer and proteins were eluted with Amylose Elution Buffer (Lysis Buffer + 10 mM maltose). MBP-R15A^325-512^ purification required an additional buffer exchange step (into Lysis Buffer) using Centri pure P1 gel filtration columns (EMP Biotech, Cat. No. CP-0110) to eliminate maltose that appeared to interfere with the dephosphorylation reactions when present at high concentrations.

<u>eIF2α^P^:</u> The N-terminal fragment of human eIF2α (1-185, with three solubilizing mutations) was purified from bacteria and phosphorylated *in vitro* using the kinase domain of PERK^2^

<u>G-actin</u> was purified from rabbit muscle according to ref. (30) as modified in ref. (10).

<u>PP1γ (7-300)</u> was purified according to ref. (13).

PP1α (7-330) was purified from BL21 *E. coli* according to ref. (27,31)

<u>PP1^N^</u> was purified from rabbit muscle according to ref. (25)

### In vitro dephosphorylation reactions

Reactions were performed at a final volume of 20 μL consisting of 5 μL of 4X solution of each component: PP1, PPP1R15A, G-actin and eIF2α^P^ (or their respective buffers). A 10X assay buffer (500 mM Tris pH 7.4, 1 M NaCl, 1 mM EDTA, 0.1% Triton X-100, 10 mM MgCl_2_) was diluted 1:10, supplemented with 1 mM DTT and used to create working solutions of PP1, PPP1R15A and eIF2α^P^ at the desired concentrations. G-actin working solutions were created using G-buffer (2 mM Tris at pH 8, 0.2 mM ATP, 0.5 mM DTT and 0.1 mM CaCl_2_). Holoenzyme components (PP1, PPP1R15A and G-actin) were combined first and substrate (eIF2α^P^) was added later to initiate the reactions, which were conducted under shaking at 500 rpm and at 30˚C for the specified time.

As a result of mixing the different proteins, the final buffer composition in the reactions was 36 mM Tris pH 7.4, 76 mM NaCl, 74 µM EDTA, 0.007 % Triton X-100, 0.7 mM MgCl_2_, 25 µM CaCl_2_, 0.05 mM ATP and 0.8 mM DTT. Additional 0.5 µM Latrunculin B ,0.4 - 3 µM MnCl_2_, 0.5% Glycerol and 50 µM TCEP were present or not depending on the proteins used on each particular reaction.

The stability test of PP1α (**Fig. S2**) was performed by preparing a fresh 240 nM solution of PP1α in the assay buffer described above. Separate aliquots were pre-incubated either at 30˚C or on ice for the specified times (30 minutes to 7 hours, see schema in **Fig. S2*A***). At termination of the preincubation, 5 μL of these pre-incubated solutions were added into 20 µL dephosphorylation reactions as described above.

Reactions performed in **Fig. 5**, included a 15 minutes pre-incubation of the enzymatic components at room temperature (before addition of substrate). Solutions of 4x PP1 (96 nM) were supplemented with either 400 μM Sephin1 (Enamine, Cat. No. EN300-195090), 400 μM Guanabenz (Sigma-Aldrich, Cat. No. D6270), 320 nM Tautomycin (Calbiochem, Cat. No. 5805551) or DMSO. Five microliters of these solutions were incubated with 5 μL PPP1R15A and 5 μL G-actin for 15 minutes before adding eIF2α^P^ substrate to initiate the reaction in a final volume of 20 µL.

**Figure 5.**
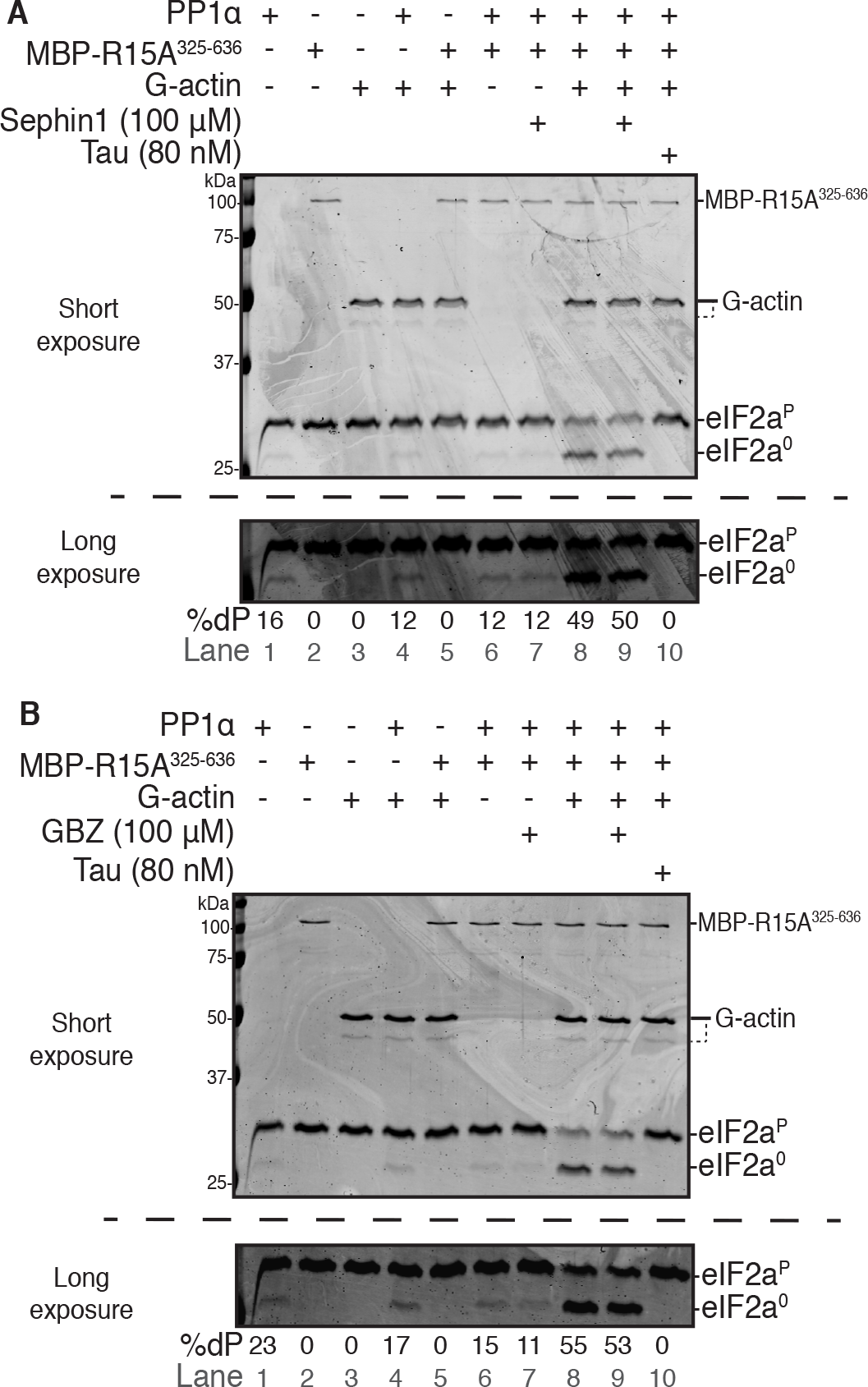
Sephin1 does not interfere with eIF2αP dephosphorylation. *A*, Coomassie-stained PhosTag-SDS-PAGE containing resolved samples from dephosphorylation reactions (20 minutes, 30˚C) in which 2 μM eIF2α^P^ was dephosphorylated by PP1α (24 nM) in presence or absence of MBP-PPP1R15A^325-636^ (60 nM) and/or G-actin (400 nM). The components were pre-incubated as specified with either Sephin1 (100 μM), Tautomycin (80 nM) or DMSO (vehicle) for 15 minutes at room temperature before being added to the reaction. The bottom panel shows a long exposure of the relevant section of the image above corresponding to the phosphorylated and non-phosphorylated forms of eIF2α. Quantification of percentage of dephosphorylation (%dP) is shown below the image. Shown is a representative experiment of three independent experiments performed. *B*, as in “*A*” but with Guanabenz (GBZ). Shown is a representative experiment of two independent experiments performed.

Reactions were terminated by addition of 10 μL of 3X Laemmli buffer supplemented with 100 mM DTT and heating the samples for 5 minutes at 70˚C. A third (10 μL) of the final volume was resolved in 12.5% PhosTag SDS gels (Wako, Cat. No. NARD AAL-107) or 15% precast Phostag SDS-PAGE gels (Alpha laboratories, Cat. No. 2614190) at 200 V for 1 hour. Gels were stained with Coomassie Instant Blue and imaged on an Odyssey imager.

ImageJ was used for band quantification and GraphPad Prism v7 was used to fit data using the ‘[Agonist] vs. response (three parameters)’ analysis function.

**Table S2** lists the number of times each experiment was performed.

## Acknowledgements

We thank Alison Schuldt and members of the Ron laboratory for their comments on the manuscript. Stefan Marciniak and Joe Chambers for useful ideas for experiments and comments on the manuscript. Robin Antrobus for mass spectrometry analysis support. Supported by a Wellcome Trust Principal Research Fellowship to D.R. (Wellcome 200848/Z/16/Z) and a Wellcome Trust Strategic Award to the Cambridge Institute for Medical Research (Wellcome 100140). M.B. was supported by a Flemish Concerted Research Action (GOA15/016). W.P. was supported by National Institute of Health R01 NS091336 and the American Diabetes Association Pathway to Stop Diabetes Grant 1-14-ACN-31. Z.C. is a PhD fellow of the Fund for Scientific Research - Flanders.

## Conflict of interest

The authors have no conflicts of interest

## Author contributions

ACC conceived the study, co-designed and conducted the experiments, interpreted the results, created the figures and co-wrote the paper. ZC co-designed the experiments, assisted with the preparation of PP1 from rabbit muscle, interpreted the results, edited the manuscript. MC expressed and purified PP1α from *E. coli*, interpreted the results, edited the manuscript. WP oversaw the expression and purification of PP1α from *E. coli*, interpreted the results, edited the manuscript. MB co-designed the experiments, oversaw the purification of PP1 from rabbit muscle, interpreted the results, edited the manuscript. DR conceived the study, co-designed the experiments, interpreted the results and co-wrote the paper

---

**Supporting Figure 1.**
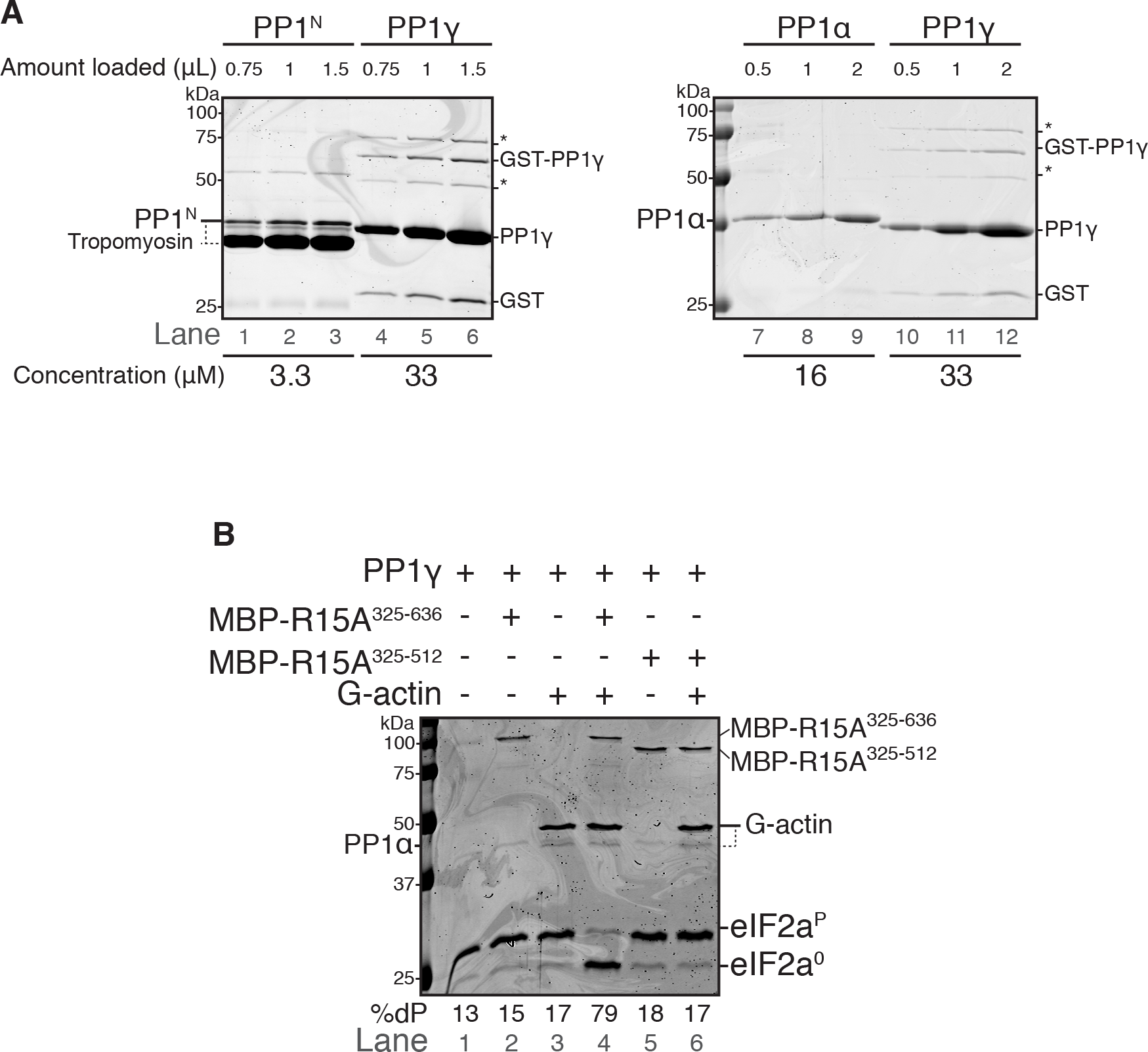
Analysis of the purity of the different sources of PP1. *A*, Coomassie-stained SDS-PAGE gels in which different amounts of PP1 sample have been resolved. The PP1^N^ preparation gave rise to two bands: a PP1 and tropomyosin band. The PP1γ preparation contained some free Glutathione S Transferase (GST) and GST-PP1 fusion protein from the purification steps, as well as other minor contaminants (*). The PP1 concentration in the different preparations is shown below the panels, calculated using PP1γ as a reference. *B*, Coomassie-stained PhosTag-SDS-PAGE containing resolved samples from dephosphorylation reactions (as in Fig. 1*A* and *B*) in which 2 μM eIF2α^P^ was dephosphorylated using bacterially-expressed PP1γ (24 nM) in presence or absence of MBP-PPP1R15A^325-636^ (50 nM), MBP-PPP1R15A^325-512^ (50 nM) and/or G-actin (400 nM) for 20 minutes at 30˚C. Quantification of percentage of dephosphorylation (%dP) is shown below the image.

**Supporting Figure 2.**
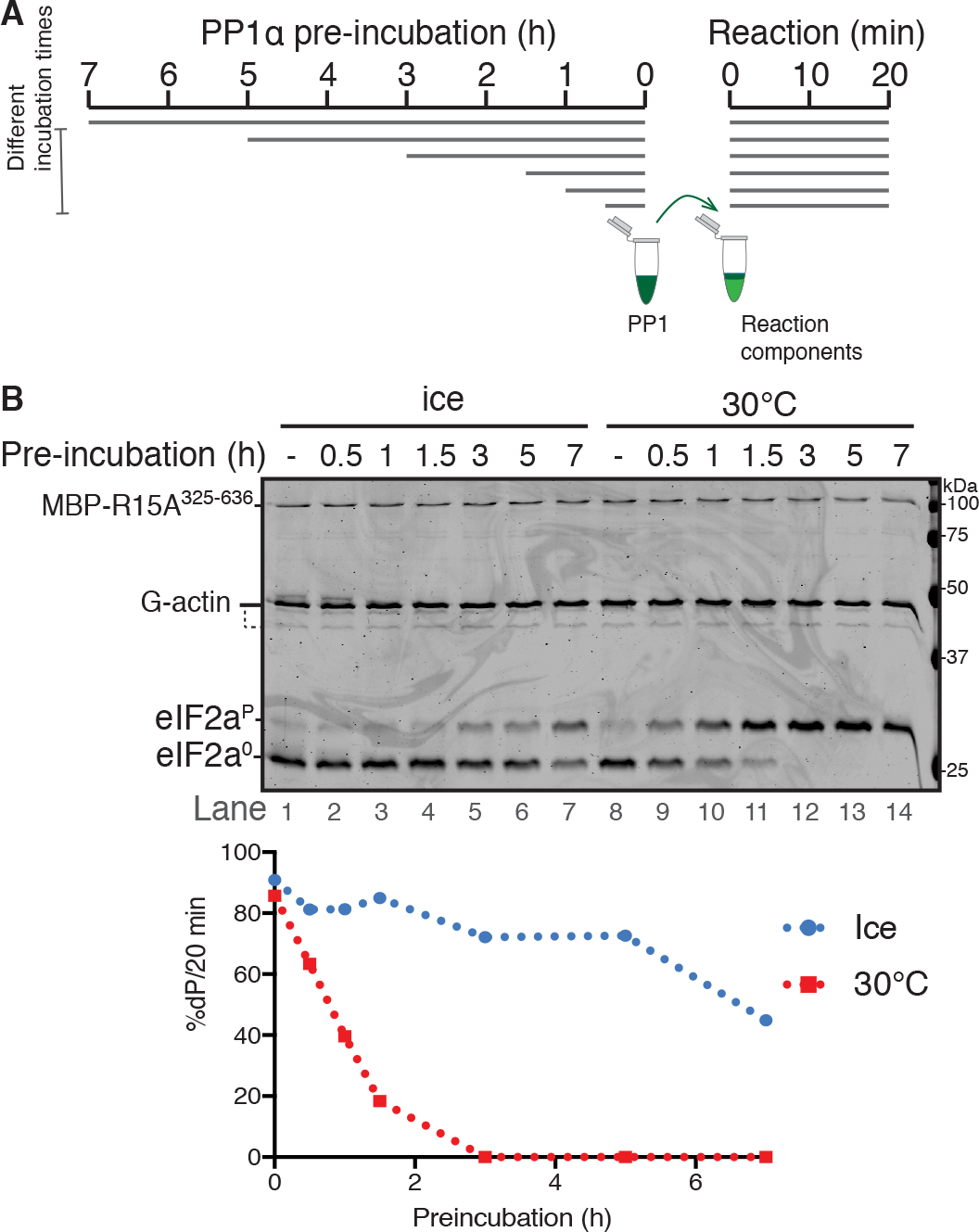
PP1α is an unstable enzyme. *A*, Schema of the experiment. Samples of PP1α (at 240 nM) were pre-incubated for the indicated period of time, either on ice, or at 30˚C, before being diluted into a eIF2α^P^ dephosphorylation reaction. *B*, <u>Upper panel</u>: Coomassie-stained PhosTag-SDS-PAGE containing samples from dephosphorylation reactions (20 minutes at 30˚C) in which 2 μM eIF2α^P^ was dephosphorylated by the pre-incubated PP1α (60 nM) in presence of MBP-PPP1R15A^325-636^ (60 nM) and G-actin (400 nM). <u>Lower panel</u>. Plot of the rate of dephosphorylation of eIF2α^P^ as a function of pre-incubation time of PP1α catalytic subunit. Data was obtained by quantification of bands of image shown above.

